# Shoot maturation strengthens FLS2-mediated resistance to *Pseudomonas syringae*

**DOI:** 10.1101/2023.02.14.528542

**Authors:** Lanxi Hu, Brian Kvitko, Li Yang

## Abstract

A temporal-spatial regulation of immunity components is essential for properly activating plant defense response. Flagellin-sensing 2 (FLS2) is a surface-localized receptor that recognizes bacterial flagellin. The immune function of FLS2 is compromised in early stages of shoot development. However, the underlying mechanism for the age-dependent FLS2 signaling is not clear. Here, we show that the reduced basal immunity of juvenile leaves against *Pseudomonas syringae* pv. tomato DC3000 is independent of FLS2. The flg22-induced marker gene expression and ROS activation were comparable in juvenile and adult stage, but callose deposition was more evident in the adult stage than that of juvenile stage. We further demonstrated that microRNA156, a master regulator of plant aging, suppressed callose deposition in juvenile leaves in response to flg22 but not the expression of *FLS2* and *FRK1 (Flg22-induced receptor-like kinase 1)*. Altogether, we revealed an intrinsic mechanism that regulates the amplitude of FLS2-mediated resistance during aging.

## Introduction

At the cell surface, some receptor kinases and receptor-like proteins act as pattern recognition receptors (PRRs) to recognize evolutionarily conserved microbe/pathogen-associated molecular patterns (M/PAMPs) and activate the Pattern-triggered immunity (PTI) (Ngou et al., 2022, Couto & Zipfel, 2016). The PTI response can be hindered by effectors, which are pathogen-derived molecules secreted into plant cells or apoplastic spaces, resulting in effector-triggered susceptibility (Jones & Dangl, 2006). Flagellin-sensing 2 (FLS2) is a receptor-like kinase localized at plasma membrane. FLS2 recognize flg22, a 22 amino acid peptide derived from bacterial flagellin, and activates PTI (Zipfel et al., 2004; Couto & Zipfel, 2016). Hallmarks of the FLS2-mediated PTI occur within minutes include changes of ion-flux at the plasma membrane, increase of cytosolic Ca^2+^ level and production of reactive oxygen species (ROS) (Seybold et al., 2014; Couto & Zipfel, 2016). Within hours, transcriptional reprogramming is induced (Li et al., 2016). In the following hours and days, deposition of callose papillae occurs to reinforce cell walls (Li et al., 2016; Couto & Zipfel, 2016). Then, synthesis of hormones amplifies immune signaling through triggering second transcriptional waves (Li et al., 2016; Couto & Zipfel, 2016). The sum of those events limit pathogen invasion and multiplication.

Misfiring of immune responses often leads to compromised growth (J. K. M. Brown, 2002, 2003; Nelson et al., 2018). Temporal-spatial regulation of activation and expression dosage of immune components are necessary to fine-tuning growth and defense (Fröschel et al., 2021; H. Wu et al., 2020; Zheng et al., 2015). FLS2-mediated signaling is developmentally regulated. In the *Arabidopsis* root, the promoter of *FLS2* was active in the differentiation zone, specifically in the vascular cylinder (Beck et al., 2014). Flg22-activated expressions of *FRK1 (FLG22-INDUCED RECEPTOR-LIKE KINASE)* and *PER5 (PEROXIDASE 57)* were confined at the cortical cell layer during lateral root formation, where root primordia pushed and damaged neighboring cortical cells (Zhou et al., 2020). FLS2 promoter activity was high in tissues that were potential bacterial entry sites, such as stomata, hydathodes, and lateral roots (Beck et al., 2014). During leaf ontogenesis, the *FLS2* transcripts were less abundant in expanding (young) leaves than that in expanded (mature) leaf, and its expression became undetectable later in senescent leaves (Klepikova et al., 2016). Zou *et al* demonstrated that the expression of *FLS2* was suppressed in early development of *Arabidopsis* cotyledon (Zou et al., 2018), leading to the higher susceptibility to bacterial pathogen *Pseudomonas syringae* (*P. syringae*) in 2-day-old seedlings than that in 6-day-old seedlings (Zou et al., 2018). In the transition from inflorescence meristem to floral meristem in *Arabidopsis, FLS2* was inhibited by a transcription factor, LEAFY (LFY). LFY suppressed the *FLS2* expression through binding to the promoter of *FLS2* (Winter et al., 2011). Hence, cauline leaves in *lfy* mutant were more resistant than that of wild type to *P. syringae* (Winter et al., 2011). Interestingly, the susceptible mutant phenotype of *fls2* to *P. syringae* was only evident in leaves generated late on the Arabidopsis shoot (Zipfel et al., 2004). To date, we are not clear about how the strength of the FLS2-mediated PTI response is regulated across different stages of shoot maturation.

In this work, we revealed that the FLS2-mediated resistance to *Pseudomonas syringae pv*. tomato DC3000 (*Pto* DC3000) (Cuppels, 1986) was increased during shoot maturation in *Arabidopsis*. We expanded the observation about *fls2* mutant phenotype in leaf from adult stage to juvenile stage. We showed that flg22-induced ROS accumulation and marker gene expression were comparable in juvenile and adult leaves. However, the level of callose deposition was higher in adult leaves than that in juvenile leaves. MiR156-regulated aging pathway showed limited impact on flg22-induced ROS production and gene activation but reduced callose deposition in fully expanded juvenile leaves. We propose that the intrinsic control of callose deposition in juvenile leaves mediates the maturation of FLS2 immune response.

## Results

### FLS2-mediated defense against Pto DC3000 was dispensable in juvenile leaves

To confirm and expand the observation of age-dependent requirement of FLS2 in resistance against *Pto* DC3000 (Zipfel et al., 2004), we measured bacterial growth in juvenile (leaves 1 & 2) and adult (leaves 13-17) leaves. Because transcriptional regulation of FLS2 occurs during leaf expansion, we used fully expanded juvenile and adult leaves to focus on the impact of shoot maturation and minimize the consequence of leaf ontogeny (Supplementary Fig. S1). The fully expanded leaves 1 & 2 and leaves 13-17 were harvested from plants grown under short-day conditions (Figure 1A and Supplementary Fig. S1). *Pto* DC3000 bacterial multiplications in two independent *fls2* alleles were not different from Col-0 juvenile leaves, while *fls2* adult leaves were more susceptible than Col-0 adults (Figure 1A-B, Supplementary Table. S1), which was consistent with a previous report (Zipfel et al., 2004). Many studies on FLS2 immune function have used bacterial surface inoculation (Zipfel et al., 2004; Zeng & He, 2010; Orosa et al., 2018). We found that juvenile *fls2* leaves were not more susceptible than Col-0 juvenile’s when *Pto* DC3000 was applied either through syringe infiltrated or spray (supplementary Table. S1). These results confirmed that the susceptibility of *fls2* relative to Col-0 wild type was age-dependent.

**Figure 1.**
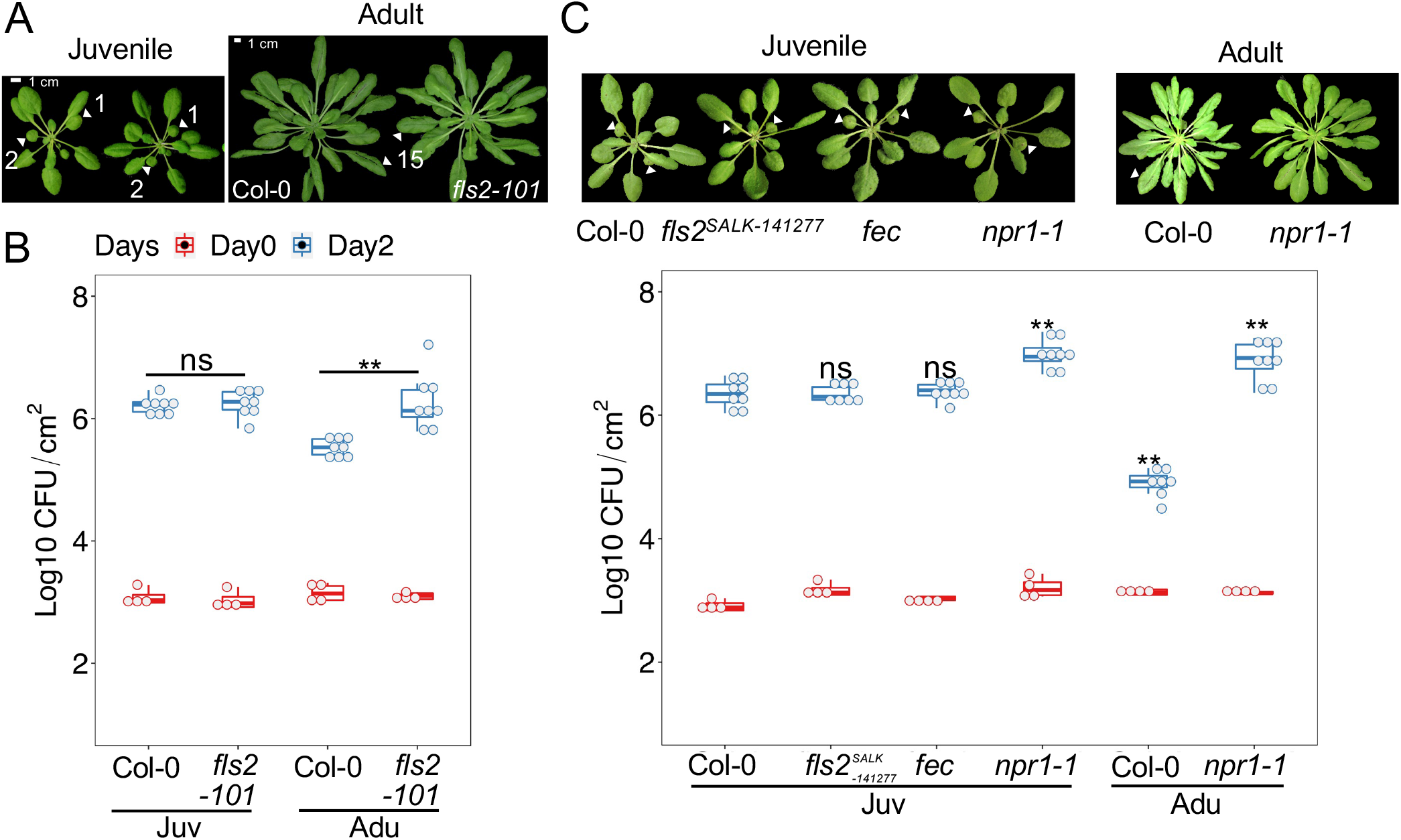
The age-related resistance to *Pto* DC3000 was abolished in *fls2* mutant. **A**, plant morphology of juvenile and adult Col-0 and *fls2-101*. **B**, *fls2* mutant was more susceptible than Col-0 in the adult stage, but not in the juvenile stage. Juv, juvenile leaves. Adu, adult leaves. Arrows indicate the leaves used for juvenile (leaves 1 and 2) and adult samples (leaf 15). **C**, *fls2 (SALK_141277), fls2/efr/cerk1* but not *npr1-1* showed bacterial growth similar to Col-0 in juvenile phase. Juv, juvenile leaves. Adu, adult leaves. Arrows indicate leaves 1 and 2 in juvenile samples; and leaves 15 in adult samples. Images of plants were taken one day before bacterial inoculation. In boxplots, each dot represents a sample that was homogenized with four leaf discs derived from four individual leaves. Red boxes indicate the initial inoculation of *Pto* DC3000 in leaves. Blue boxes indicate bacterial growth in leaves two days post-inoculation (dpi). The bacteria growth was estimated by counting bacterial colony forming unit/cm^2^ (CFU/cm^2^). * P < 0.05, ** P < 0.01, student t-test. The data are representative of three experimental repeats that were performed with similar results.

We first speculated that the lack of phenotype in *fls2* juvenile leaves was due to a redundancy of FLS2-mediated defense with other PTI pathways. We tested bacterial growth in *fec*, the triple mutant of loss-of-function *FLS2/EFR/CERK1* immune receptors (Gimenez-Ibanez, Ntoukakis, et al., 2009). Like *fls2*, leaves 1-2 from *fec* mutant showed the same level of susceptibility to *Pto* DC3000 as that of Col-0 (Figure 1C), arguing against the possibility of the functional redundancy among the three signaling pathways. The result implied that a downstream of *FLS2/EFR/CERK1* was compromised in juvenile leaves. The *npr1-1* mutant, defective in salicylic acid perception (Cao et al., 1997), showed increased bacterial propagation in juvenile leaves (Figure 1C), suggesting the bacterial load was not constrained by the physiology of juvenile leaves. This data suggests that FLS2-mediated defense signaling to *Pto* DC3000 is hindered in juvenile leaves.

### Inefficiency of FLS2-mediated resistance in juvenile leaves was not due to the functions of effectors and coronatine

*Pto* DC3000 delivers effector proteins and toxin coronatine to suppresses FLS2-mediated defense (Gimenez-Ibanez, Hann, et al., 2009; Shan et al., 2008; Xiang et al., 2008; Zeng & He, 2010). We sought to assess if the inefficiency of the FLS2 signaling in juvenile leaves were due to those virulent factors. We separately inoculated juvenile leaves with three *Pto* DC3000 mutant strains: *hrcC-*, defective in effector delivery through Type III Secretion System (T3SS) (Yuan & He, 1996); *cor-*, the Δ*cfl*Δ*cfa9 (cfa, coronafacic acid; cfl, cfa ligase)* (Bender et al., 1993) or the Δ*cfa6* mutant (Zeng et al., 2011) that is deficient in the phytotoxin coronatine biosynthesis, and the *hrcC-/cor-* double mutant (Worley et al., 2013). In *fls2-101* juvenile leaves the *hrcC-, cor-* and *hrcC-/cor-* strains reached similar respective loads to what was observed in the Col-0 juvenile leaves (Figure 2A-C, supplementary Table. S1). Neither the high inoculum (OD600=0.1) nor the additional period of growth made a difference for *hrcC-/cor-* growth in Col-0 and *fls2* juvenile leaves (Figure 2C, supplementary Table. S1). When compared with adult leaves, juvenile leaves were more susceptible to those mutated strains, displaying an age-dependent defense response (supplementary Table. S1). Thus, the evidence supports that host factors or additional pathogen virulence factors constrained the FLS2-defense in juvenile leaves.

**Figure 2.**
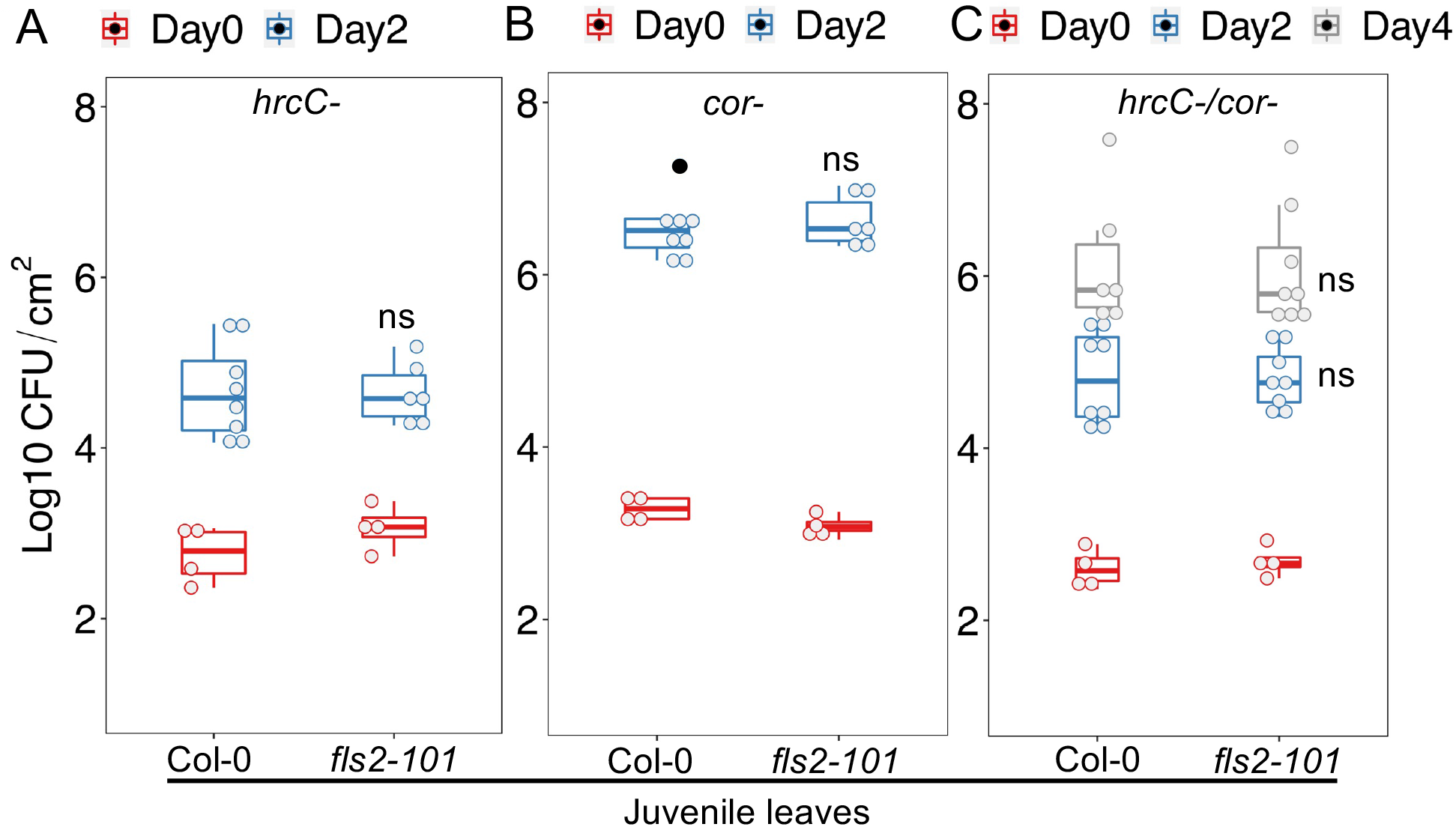
The growth of *Pto* mutants was comparable in juvenile leaves of *fls2* and Col-0. **A**, comparable bacterial growth of *hrcC-* in juvenile Col-0 and *fls2*. Each dot represents a sample containing 4 leaf discs. **B**, a comparable growth of *cor-* in juvenile leaves of Col-0 and *fls2*. **C**, the similar growth of *hrcC-/cor-* in juvenile Col-0 and *fls2*. Red boxes indicate the initial inoculation of *Pto* DC3000 and *hrcC-* in leaves. Blue boxes indicate bacterial growth in leaves two days postinoculation (dpi). Grey boxes indicate bacterial growth four days post-inoculation (dpi). The bacteria growth was estimated by counting bacterial colony forming unit/cm^2^ (CFU/cm^2^). The outliers indicated by black dots were determined by mean±two standard deviations (sds) in R. * P < 0.05, ** P < 0.01, student t-test. The data are representative from at least three experimental repeats that were performed with similar results.

### A subset of FLS2-mediated defense response was compromised in juvenile leaves

We next investigated host factors that might limit FLS2 function in juvenile leaves. To check whether *FLS2* is differentially expressed or induced between juvenile and adult leaves, we infiltrated a series dilution of flg22 in those leaves. Both basal level and flg22-induced expression of *FLS2* were comparable in juvenile and adult leaves. Expression of a PTI marker *FRK1 (Flg22-induced Receptor-like Kinase 1)* was also indistinguishable between those two stages (Figure 3A-B). Likewise, the flg22-induced production of reactive oxygen species (ROS) was not impaired in the juvenile leaves, nor was the total ROS accumulation (Figure 3C).

**Figure 3.**
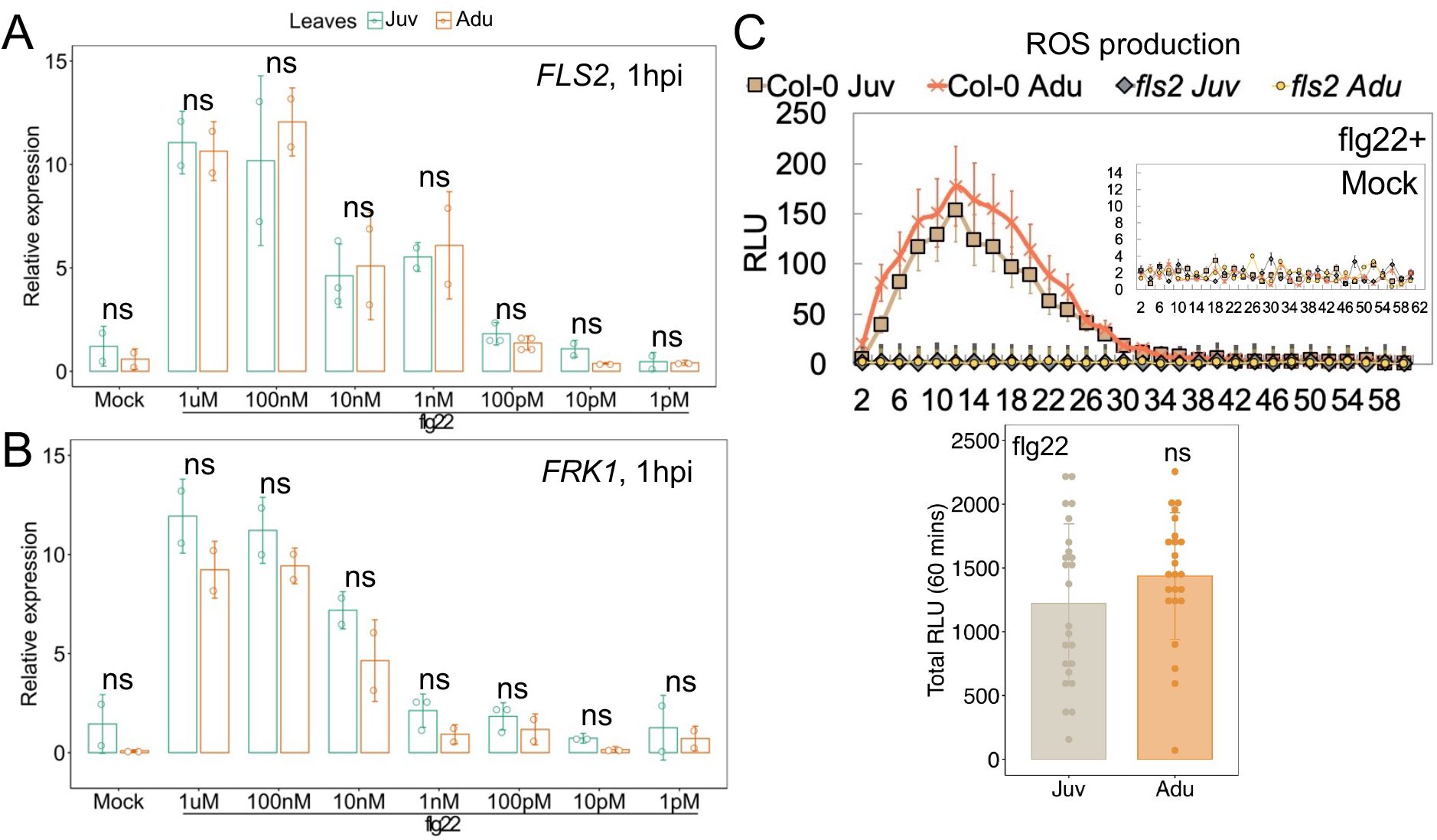
The early FLS2 immune responses were independent of shoot maturation. **A-B**, the transcript quantity of *FLS2* and *FRK1* was not temporally regulated. Juv, juvenile leaves. Adu, adult leaves. 1 *µ*M flg22-treated juvenile and adult leaf samples were harvested at 1-hour post-infiltration (hpi). Mock, 1 *µ*M DMSO in ddH_2_O. Each dot represents a technical repeat. Error bars stand for standard deviation (±SD). * P < 0.05, ** P < 0.01, student t-test. The data are representative from three experimental repeats that were performed with similar results. **C**, ROS induction and accumulation within an hour reached to the similar amplitude in juvenile and adult Col-0 leaves. See details of mock prep in methods. Flg22: 50 nM. Each symbol in the curve plot stands for an average value of at least 24 individual leaf discs. Each dot in the bar plot represents the sum of values of a single leaf disc evaluated within 60 minutes. RLU, relative light unit.

Flg22 induced callose deposition in both juvenile and adult leaves, which was dependent on FLS2 (Figure 4A-B). However, flg22-triggered callose deposition was weaker in juvenile leaves than that in adult leaves (Figure 4A-B). Next, we assessed the flg22 priming effect between the two phases of leaves. The pretreatment of flg22 protected leaves against *Pto* DC3000 in juvenile leaf of Col-0 (Figure 4C, supplementary Table. S1). Being consistent with enhanced callose deposition in the adult leaves (Figure 4A-B), bacteria multiplication was still lower in adult leaves than those in juvenile leaves after flg22 treatment (Supplementary Table. S1). Those results indicated that low callose deposition correlates with limited FLS2-mediated defense in juvenile leaves.

**Figure 4.**
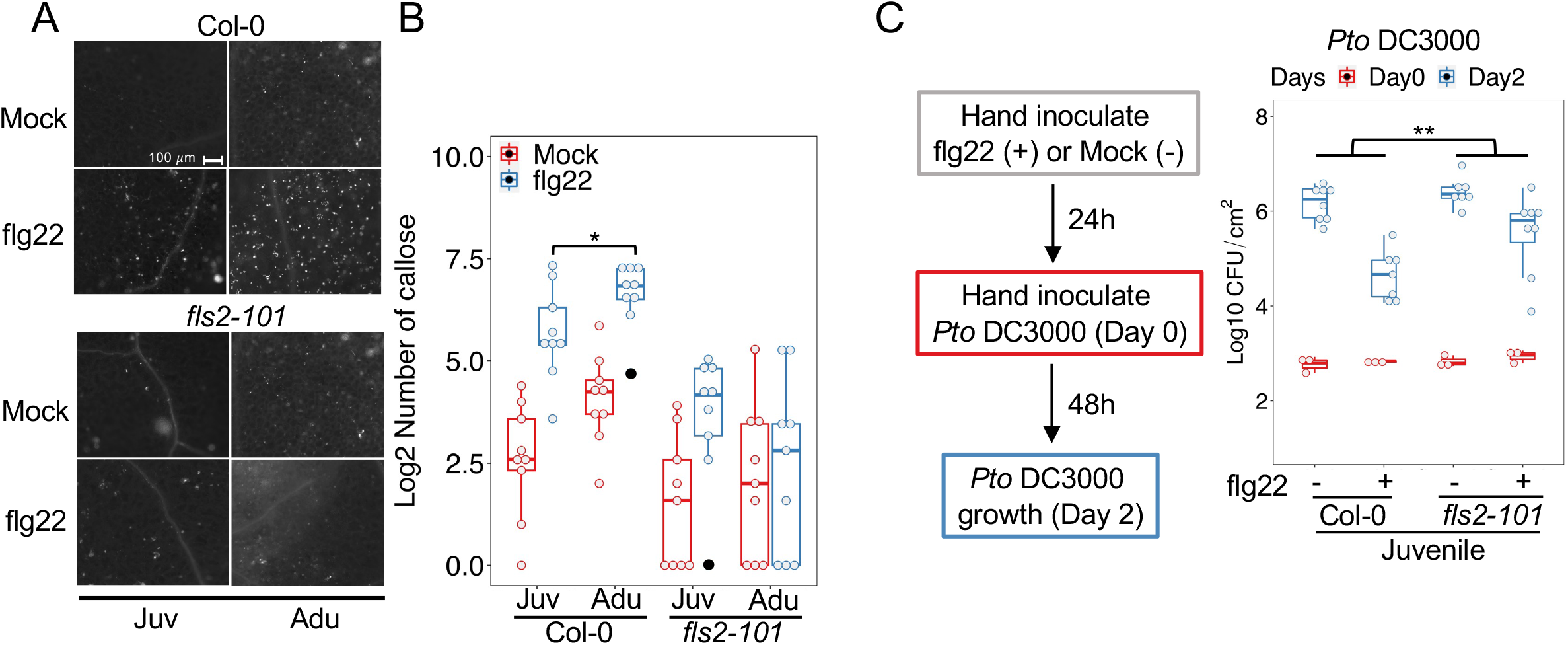
Perception of flg22 activated a weak callose deposition in juvenile leaves. **A**, visualization of callose deposited in juvenile and adult leaves of Col-0 24 hours post flg22 treatment. Flg22: 1 *µ*M. **B**, quantification of the callose deposition depicted in Fig 4A. Juv, juvenile leaves. Adu, adult leaves. The outliers indicated by black dots were determined by checking the statistical model in R. The treatment effect within each genotype or age was determined by student t test, P < 0.05, ** P < 0.01. **C**, Flg22-priming protected juvenile Col-0 compared with that in *fls2*. Flg22 was infiltrated in leaves 24h prior to inoculation of *Pto* DC3000. Left panel, a diagram of procedures of flg22-protection assay. Right panel, each dot represents a sample with four leaf discs. Red boxes indicate the initial inoculation of *Pto* DC3000 in leaves. Blue boxes indicate bacterial growth in leaves two days post-inoculation (dpi). The bacteria growth was estimated by counting bacterial colony forming unit/cm^2^ (CFU/cm^2^). The outliers indicated by black dots were determined by mean ± two standard deviations (sds) in R. Emmeans in R (Searle et al., 1980) was used in Fig 4C to determine the genotype effect on bacterial growth between treatments. The data are representative from three experimental repeats that were performed with similar results.

### The flg22-triggered callose deposition was weakened by high miR156 level in juvenile phase

MicroRNA156 (miR156) is a master regulator of the shoot maturation (Poethig, 2013). miR156 accumulates highly in plants to maintain juvenile phase (J.-W. Wang et al., 2008; G. Wu et al., 2009; G. Wu & Poethig, 2006). The temporal decline of miR156 expression allows the transition to adult phase (J.-W. Wang et al., 2008; G. Wu et al., 2009; G. Wu & Poethig, 2006). Knocking down the function of miR156 by target mimicry, *MIM156*, leads to precocious adult traits on leaves 1 and 2 (Franco-Zorrilla et al., 2007; Wu et al., 2009; Wu & Poethig, 2006). We sought to test whether miR156 controls the age dependent FLS2 immune signaling. First, we examined whether the miR156 influences the flg22-induced defense outputs using estradiol-inducible transgenic lines with either knocking down of miR156 function, *est::MIM156* (*inMIM156*) (Brand et al., 2006; He et al., 2018), or overexpressing *MIR156A, est::MIR156A* (*inMIR156*) (Brand et al., 2006). The temporary expression of *MIM156* or *MIR156A* through estradiol induction minimized the impact of morphological differences caused by the altered miR156 activity or transcriptional expression. Neither *inMIM156* juvenile leaves nor adult leaves of *inMIR156* changed ROS activities, *FLS2* or *FRK1* expression, at resting state or upon FLS2-defense activation (Figure 5A-C). These observations agreed with those results that ROS activities, *FLS2* or *FRK1* expression were not differentially regulated in the juvenile and adult leaves (Figure 3).

**Figure 5.**
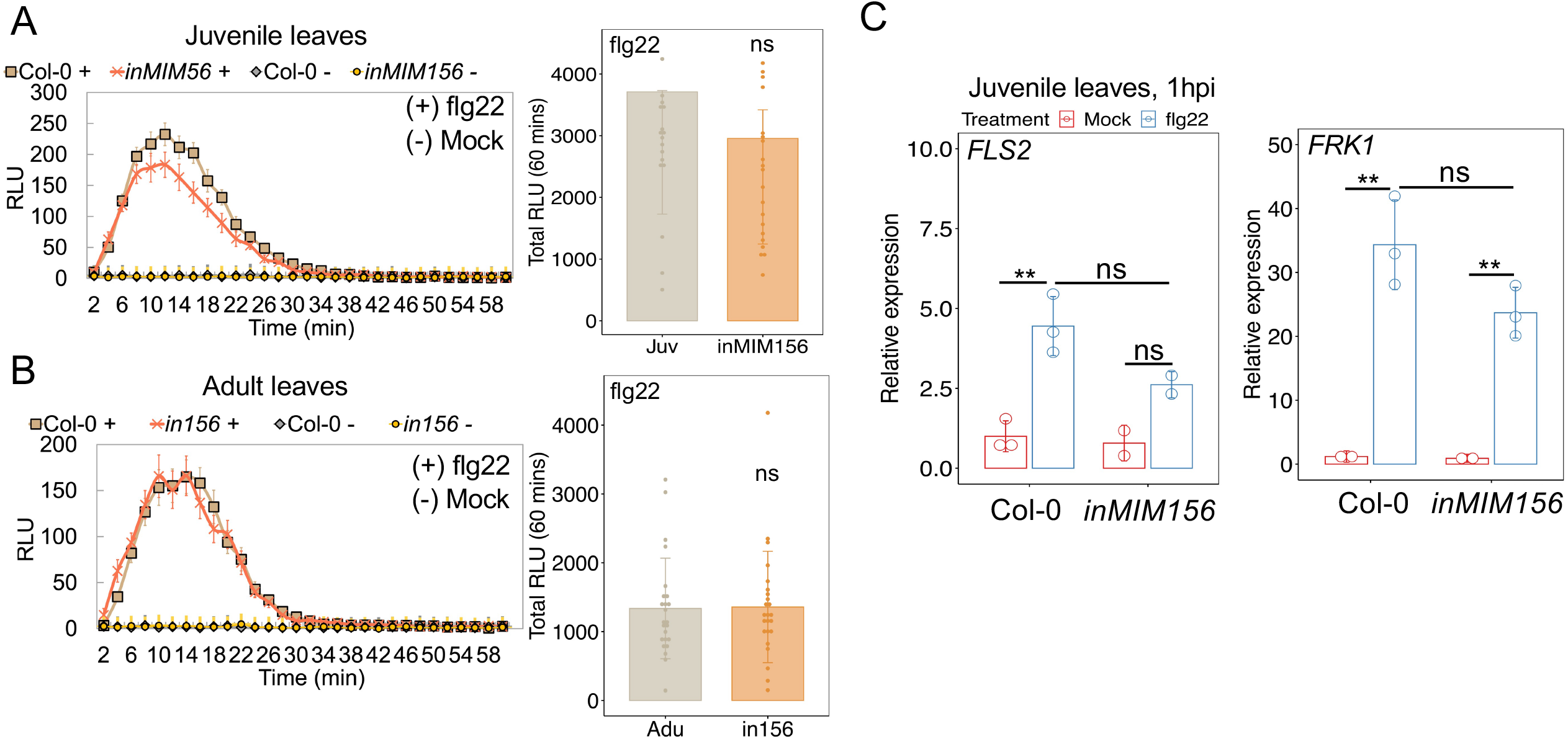
Early defense signaling in juvenile leaves was independent of miR156. **A**, Similar levels of ROS induction and total accumulation of juvenile Col-0 and *inMIM156* leaf 1-2 within an hour. **B**, Similar levels of ROS activation and accumulation of adult Col-0 and *in156* leaf 13-17. Treatments and the meaning of symbols were the same as described in Fig 3C. **C**, *FLS2* and *FRK1* expressions were not higher in leaf 1-2 of *inMIM156* than juvenile leaves of Col-0. Mock treatment refers to leaves infiltrated with 1 *µ*M DMSO. Flg22, 1 *µ*M. Samples were harvested at 1 hpi of treatments. Each dot represents a technical repeat. Error bars represent standard deviation (±SD). P < 0.05, ** P < 0.01, student t-test. The Data are representative from three experimental repeats that were performed with similar results.

Notably, induced *MIM156* led to an increased number of deposited callose at cell wall in juvenile leaves after flg22 treatment (Figure 6A-B), suggesting that the high accumulation of miR156 compromised this defense output of FLS2 signaling. However, the callose deposition phenotype in *inMIM156* juvenile leaves was still significantly lower than that in wild type adult leaves (Figure 6A-B), indicating additional age-dependent mechanisms contribute to high callose deposition potential in the adult stage or suppress which in the juvenile stage. We subsequently crossed 35S::*MIM156* (Franco-Zorrilla et al., 2007; Wu et al., 2009; Wu & Poethig, 2006) into *fls2-101* mutants to endow adult feature to the leaves 1&2 of *fls2* (Figure 6C). Consistent with our previous observation (Hu et al., 2022) that knocking down miR156 function enhanced resistance against *Pto* DC3000, leaves 1&2 in *MIM156* plants showed low bacterial multiplication. Loss of *fls2* in the *MIM156* background did not increase disease susceptibility in leaves 1&2 (Figure 6D). The similar disease phenotype of *MIM156* and *fls2/MIM156* indicates that miR156 acts downstream or in parallel with FLS2. We conclude that miR156 suppressed flg22-induced callose deposition, which together with miR156-independent factors restrict the output of FLS2 signaling in early shoot development.

**Figure 6.**
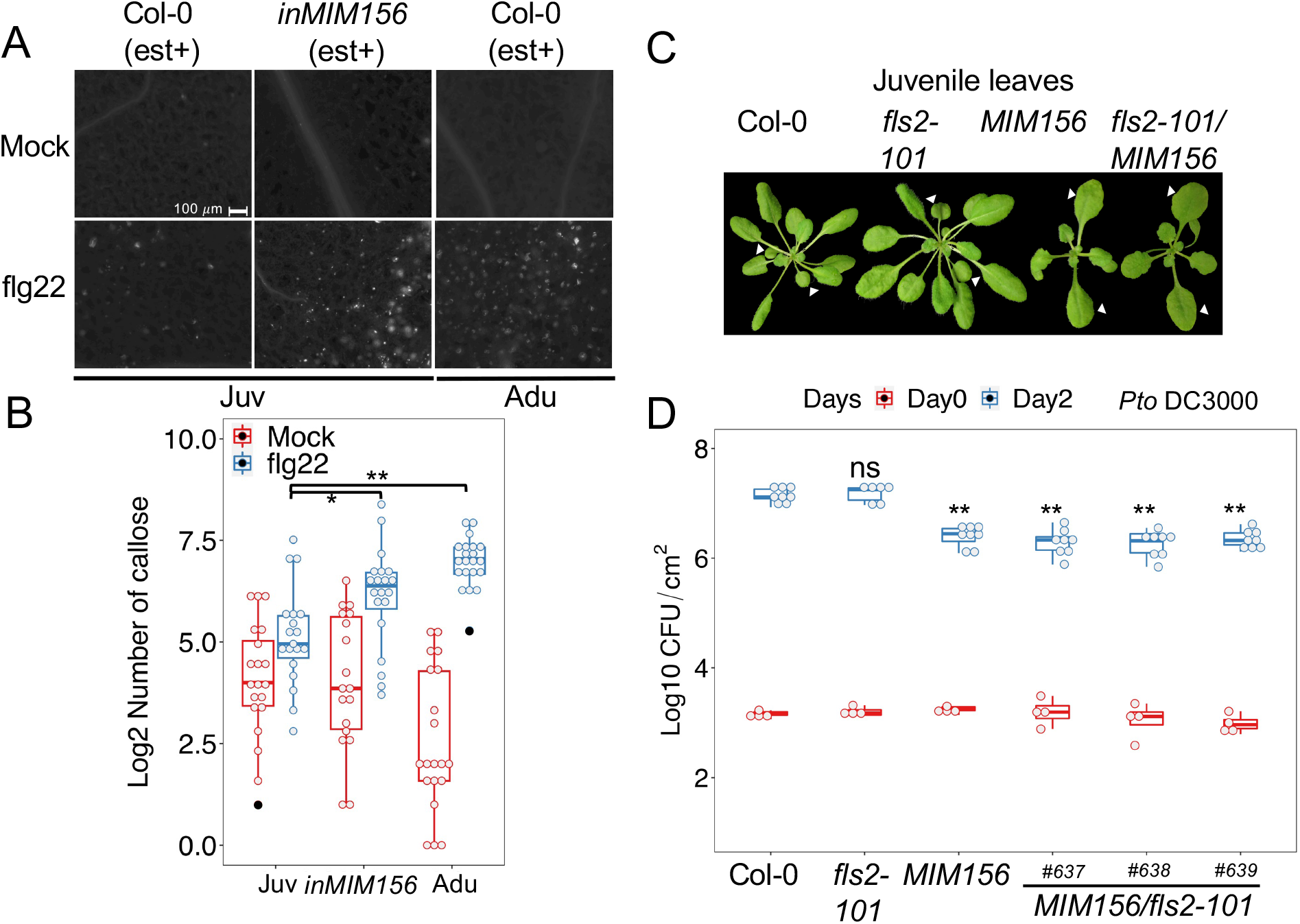
The low level of miR156 allows enhanced callose deposition and disease resistance. **A**, flg22 induced less callose deposition in juvenile Col-0 leaves than those from *inMIM156* and adult Col-0. **B**, Quantification of the callose deposition in Fig 6A. Juv, juvenile leaves. Adu, adult leaves. The black square indicated numbers of callose calculated from wounded tissue. The outliers indicated by black dots were determined by mean ± two standard deviations (sds) in R. The student t test was used between treatments within a single genotype/age, P < 0.05, ** P < 0.01. Emmeans in R (Searle et al., 1980) was used to determine genotype/age effects on the accumulation of flg22-induced callose. **C**, plant morphology of *fls2/MIM156* relative to *fls2* and *MIM156*. The photo was taken one day before the bacterial infiltration. **D**, Bacterial growth in leaves 1 and 2 from *MIM156* and *MIM156*/*fls2*. Each dot represents a sample. Red boxes indicate the initial inoculation of *Pto* DC3000 in leaves. Blue boxes indicate bacterial growth in leaves two days post-inoculation (dpi). The bacteria growth was estimated by counting bacterial colony forming unit/cm^2^ (CFU/cm^2^). P < 0.05, ** P < 0.01, student t-test. The Data are representative from three experimental repeats with similar results.

## Discussion

In this research, we reported FLS2 contributed to age-related resistance to *Pto* DC3000 in adult leaves in *Arabidopsis*. We measured FLS2 immune signaling cascade including the flg22-induced ROS burst, the level of *FLS2* and *FRK1* transcripts and callose deposition. We discovered that the callose deposition phenotype was dampened in juvenile leaves. In support of that, the pre-treatment of flg22 decreased the bacterial load in adult leaves when compared with what was in flg22-treated juveniles. Furthermore, high accumulation of miR156 in the juvenile stage hindered flg22-triggered callose deposition, with potentially additional factors together led to the inefficiency of FLS2 resistance (Figure 7). Our work provided a temporal regulation on a subset of FLS2-mediated PTI responses, which associates with age-dependent defense response.

**Figure 7.**
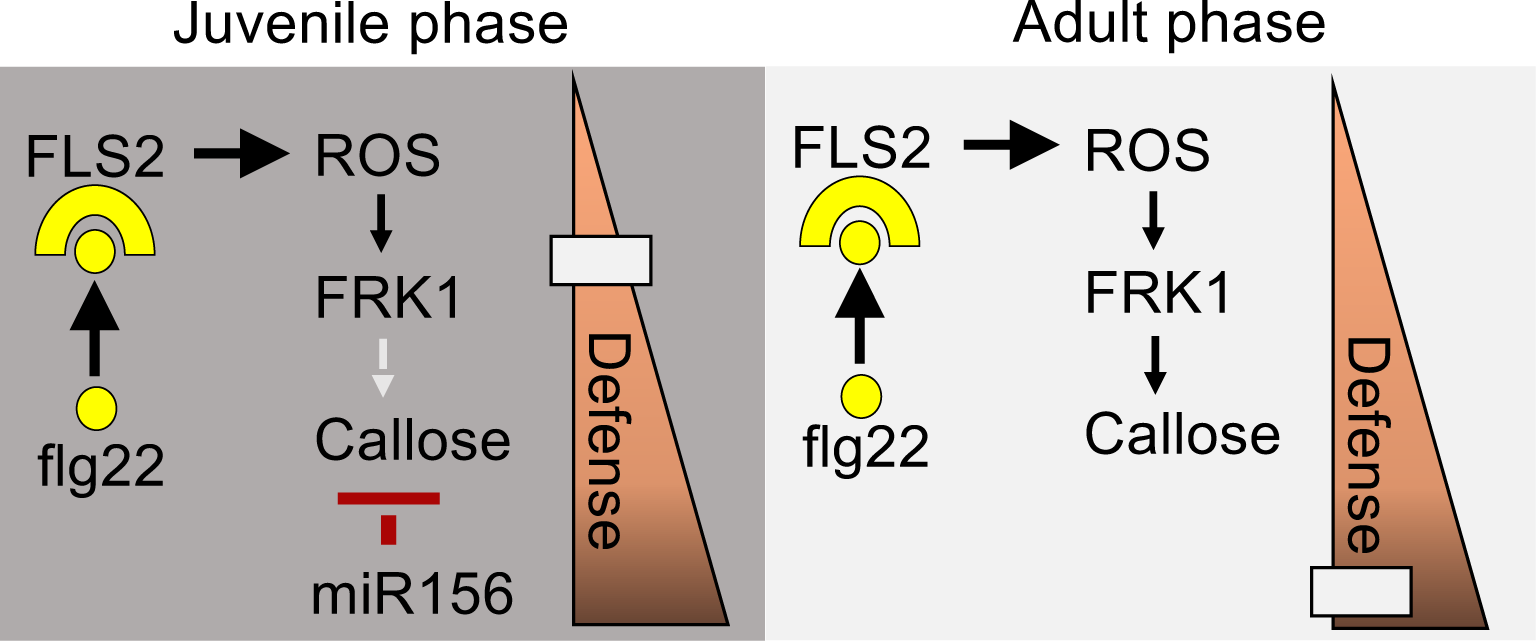
A diagram for the proposed model. Flg22 induces FLS2-mediated PTI in juvenile and adult phase. Callose deposition, but not ROS signaling, accumulation or early marker activation, was compromised in the juvenile phase. Temporal reduction of miR156 contributed to the increase of callose deposition in adult stage. Additional host factors may limit FLS2-mediated PTI or promote which in the juvenile stage.

The change of immune strength during vegetative phase change has been documented in various plants. The overexpressing enhancer mutant of the miR156, *Corngrass 1*, led to extended juvenile phase and increased susceptibility to common rust (*Puccinia sorghi Schw*) and European corn borer (*Ostrinia nubilalis Hubner*) (Abedon & Tracy, 1996). Silencing miR156 in *MIM156* mutant accelerated adult development in rice leaf and increased resistance to brown planthopper (Ge et al., 2018). In other cases, a weak allele of *Hm1, Hm1*^*A*^, conferred adult resistance to northern leaf spot in maize (Marla et al., 2018); Cf-9B-mediated resistance to leaf mold in tomato was incremental over the vegetative-reproductive transition (Panter et al., 2002). Neither *Hm1*^*A*^ nor *Cf-9B* changed transcriptional expression within those developmental transitions (Marla et al., 2018; Panter et al., 2002). The alternative splicing of *Cf-9B* was alluded to play a role in the onset of mature resistance (Panter et al., 2002). Whether miR156 participates in those age-related resistances remains to be seen.

Callose deposition to the plant cell wall contributes to defense against pathogen invasion, especially for preventing the penetration of fungal hyphae (An, Ehlers, et al., 2006; An, Huckelhoven, et al., 2006; Nielsen et al., 2012). Many Gram-negative bacterial pathogens require T3SS-secreted effectors to suppress callose deposition. T3SS-deficient strains of *Pto* DC3000 and *Xanthomonas euvesicatoria* 85-10 induced abundant callose deposition, thickening cell walls and triggering papilla formation (Bestwick et al., 1995; I. Brown, 1995; I. Brown et al., 1998). The pathogen-induced papillae are a cell wall apposition for plants to deliver defense components, such as phytotoxin, callose and ROS (Meyer et al., 2009; Y. Wang et al., 2021). AvrPto effector derived from *Pto* DC3000 repressed callose papillae deposition and enabled considerable multiplication of the T3SS-deficient strain of *Pto* (*hrp* mutant, Hauck et al., 2003). A decrease of papillae was observed in miR156-overexpressing rice plants (Xie et al., 2012), However, overexpression of miR156 in switchgrass increased lignin content and decreased susceptibility to fungal rust (*Puccinia emaculata*), while the plant became highly susceptible to Bipolaris species (Baxter et al., 2018). As a part of constitutive defense, the biosynthesis of wax and cuticle was upregulated in adult than that of juvenile Col-0 leaves (Hu et al., 2022). In maize and *Arabidopsis*, miR156 regulates cell wall composition during the vegetative phase change (li et al., 2019; Strable et al., 2008; Vega et al., 2002). Investigating the modulation of miR156 in cell wall-defense in the absence and presence of pathogens would be of next interests.

The magnitude of flg22-triggered ROS signaling and the *FRK1* expression are maintained during the vegetative transition. Since miR156 functions in the absence of pathogen attack, it is plausible that the high level of miR156 inhibited basal activities of defense, which then attenuated a subsector of FLS2-mediate PTI. To which extent that a high level of defense gene expression at steady state contribute to defense activation is unclear in plants. Future research is required to determine the regulatory components and their modes of action pre- and post-infection in the age-related resistance. Quantitatively analyze different sectors of PTI responses in the distinct stage of development would be informative to understand the aging-immunity crosstalk at molecular levels and strategize disease managements accordingly. The age-related resistance can be maintained for more than 30 years when crops grow substantially in the region exposed with multiple races of pathogens (Roland F. Line & Xianming Chen, 1995). The adult resistance in wheat is conferred by different nonclassical R genes, which cannot be overcame by a single genetic variation (Ellis et al., 2014; Moore et al., 2015; Schwessinger, 2017). Limiting the population size of multiple pathogens without demanding a fast genomic expansion of resistance gene can make the age-related resistance an efficient tactic for plants to stay resilient under turbulent environmental conditions.

---

**Supplementary Fig S1.**
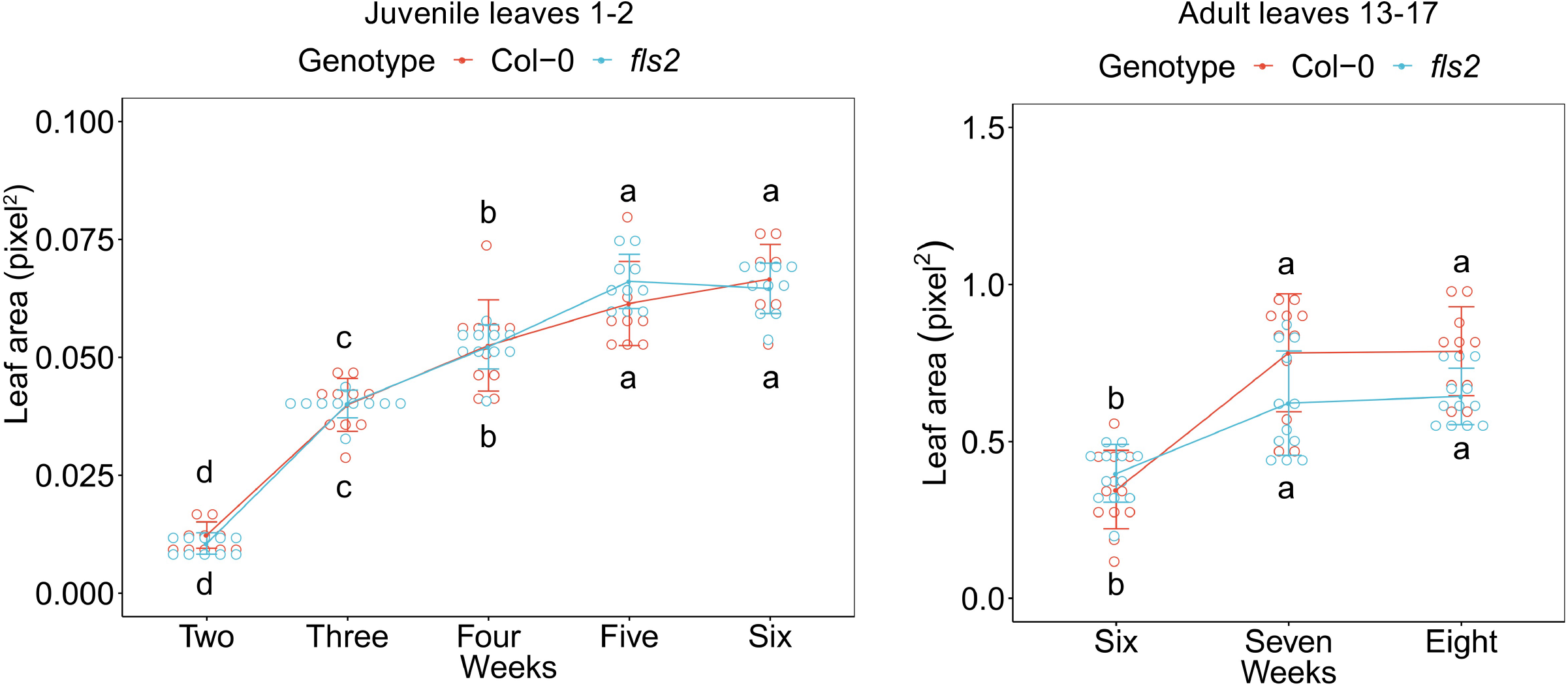
*Fls2* mutant had comparable leaf expansion rates as that of Col-0. In the curve-dot plots, each dot stands for a single leaf. Tukey test was applied for statistics.

**Supplementary Table1. Experimental repeats of bacterial growth.**

**Supplementary Table2. Plants, bacteria strains and primers used in this study**.

**Supplementary information. Statistics in R**.

## Materials and Methods

### Plant material and growth conditions

Planting and growing conditions (9 hours light/15 hours darkness) are the same as described in (Hu et al., 2022). All juvenile leaves of Col-0 and leaf 1 and 2 of mutants that were used in this work coming from 4-5 weeks old plants in soil. All adult leaves of Col-0 and leaves at the similar positions of mutants were from 6-7 weeks old plants in soil. *Fls2-101, fls2* (SALK_141277), and *npr1-1* mutant were obtained from the Arabidopsis Biological Resource Center (Ohio State University, Columbus, OH). The cross of *fls2-101* and *35S::MIM156* were generated from this research, and genotyped with primers in supplementary Table. S2. Progenies in F3 and F4 of *fls2/MIM156* were used for phenotypic tests.

### Bacterial growth assay

*Pto* DC3000 and *Pto* mutants were grown at 28 °C on King’s B medium (40 g l^-1^ proteose peptone 3, 20 g l^-1^ glycerol, 15 g l^-1^ agar) with Rifamycin (final concentration 100 *µ*g/mL). An overnight fresh plate culture of each bacterial strain was prepared. For needleless syringe infiltration, bacteria were scraped off the plate and resuspended in 10 mM MgCl_2_ (OD=0.1 and then with 500 times of dilution). For spraying the DC3000 strain, bacterial cell suspension (OD=0.1) was prepared in 10 mM MgCl_2_ with 0.01% Silwet L-77. After inoculation, leaves from four individual juvenile or four adult plants were collected separately and homogenized (homogenizer, OMNI International) in one biological replicate. The homogenized samples were then loaded on fresh King’s B plate with serial dilutions. Two days after loading, single colonies were counted manually. Usually three-four biological replicates per genotype/age per treatment were collected at Day0, and six-eight biological replicates were used for Day2 and Day4.

### Flg22-induced protection assay

1*µ*M of flg22 or mock (1 *µ*M DMSO in ddH2O) were infiltrated in leaves with needleless syringe 24 hour prior to the bacteria infection. Infiltration of *Pst* DC3000 and sampling are the same as described in the bacterial growth assay.

### Gene expression analysis

To detect flg22-induced activation of *FLS2* and *FRK1*, 1 *µ*M, 100 nM, 10 nM, 1nM, 100 pM, 10 pM and 1 pM of flg22 (GenScript) were infiltrated in leaves with a needleless syringe. Since flg22 was dissolved in DMSO, 1 *µ*M DMSO in ddH2O was used as mock. Samples were harvested at 1 hpi. Twenty juvenile leaves or 20 adult leaf discs from at least three individual plants were collected and homogenized (homogenizer, OMNI International) as one biological replicate. One to two biological repeats were used in a single experiment. Total RNAs were extracted using E.Z.N.A. Total RNA kit (Omega BIO-TEK) and reversely transcribed with GoScript reverse transcriptase (Promega). The qPCR was performed using SYBR Green master mix (Applied Biosystems) in the QuantStudio 1 Real-Time PCR system (Applied Biosystems). qPCR conditions were the same as described in Hu et al., 2022. Reference genes were *TUB2* (*AT5G62690*) and *SAND* (*AT2G28390*). Primers of qPCR were shown in supplementary Table. S2.

### Callose deposition assay

The protocol was adapted from (Yu et al., 2019). 1 *µ*M flg22 or mock (1 *µ*M DMSO in ddH2O) was hand-infiltrated in juvenile and adult leaves of Col-0, *fls2-101* as well as the leaf 1 and 2 of *inMIM156*. Nine to twenty leaves from 8 plants (sample size varied between experimental repeats but was kept similar in an experiment between groups) were harvested 24h post the treatments. Leaf discs were collected from the same position among adult leaves using a corer that has comparable sampling area with the size of juvenile leaves. The samples were fixed with FAA solution (10% formaldehyde, 5% acetic acid and 50% ethanol) via vacuum infiltration followed by an overnight incubation under 37 °C. Using 95% ethanol to incubate the samples 24h to clear chlorophyII pigment. Rinse the samples with 75% ethanol each day over 2-3 days, until leaves became transparent. Rinse samples once with ddH_2_O. Stain the samples for 30 min with 0.01% aniline blue solution (150 mM KH_2_PO_4_, pH 9.5). At the same day after the staining, imaging all the samples using fluorescent microscope (Zeiss). The number of callose was quantified using Fiji software.

### ROS assay

Following the protocol derived from (Sang & Macho, 2017), Biopsy punch with plunger (4 mm diameter; Miltex, USA) was used to collect leaf disc from juvenile and adult leaves of Col-0, *fls2-101, inMIM156* (leaf 1-2) and *in156* (leaves at the same position with that of adult Col-0). At least 24 leaf discs per phenotype per treatment was used for each experiment. Each leaf disc was placed in an opaque 96-well plate (OptiPlate™-96, Perkin Elmer) and submerged in 100 *µ*L of ddH_2_O overnight. After 16-18 hrs, the water in the plate was replaced by a master mix with the same volume in each well. The total of 10 mL master mix solution was made of 10 *µ*L of Luminol (Sigma) from 100 mM stock solution in DMSO, 10 *µ*L of Horseradish Peroxidase (Sigma) from 20 mg/mL stock solution in ddH_2_O, 5 *µ*L of flg22 peptide (100 *µ*M working stock) and supplemented with ddH_2_O. To make the 100 *µ*M of flg22 working stock, 10 *µ*L of elicitor at 10 mM (dissolved in DMSO) was added in 990 *µ*L of ddH_2_O. As a negative control, the same master mix solution was made with flg22 replaced by DMSO. The H_2_O_2_-reacted luminescence was detected under luminometer (Molecular Devices, SpectraMax iD3) from 2 min up to an hour. The data was collected and analyzed in Excel software.

### Construction of inducible MIR156A

The estradiol-inducible *MIR156A* line (*inmiR156*) was constructed using a Gateway compatible version of the XVE system (Brand et al., 2006). The *MIR156A* stem loop sequence was cloned into pMDC160. Homozygous transgenic line was crossed to plants containing pMDC150-35S (Brand et al., 2006) and selected for double homozygous. The working concentration of 20 *µ*M 17-ß-estradiol (dissolved in DMSO and diluted with ddH2O plus 0.01% Silwet 77) was sprayed onto leaves at 12 hours before collecting leaf samples for ROS assay. For qPCR assay, estradiol was treated 24 hours before sample collections.

## Data visualization and statistics

The data visual was done using Microsoft Excel and RStudio 2022.07.1+554 (RStudio Team, 2022) with package ggplot2 3.3.6/ggpubr0.4.0 (Wickham, 2016), package magrittr 2.0.3 (Stefan Milton Bach & Hadley Wickham, 2022), and package RColorBrewer 1.1-3 (Brewer et al., 2003). The student t test was performed in the T test calculator, GraphPad, https://www.graphpad.com/quickcalcs/ttest1.cfm. Emmeans package 1.8.1-1 was conducted in RStudio (Searle et al., 1980).

## Acknowledgement

We thank undergraduate students assisting experiments in the Yang lab. We appreciated the critical feedback of this research from Chang Hyun Khang and Shavannor M. Smith. We thank Paul Severns to provide advice for statistics. This project is supported by NIH R35GM143067 to Y.L.

