## Supplementary material for "Shoot maturation strengthens FLS2-mediated resistance to *Pseudomonas syringae*": sup_information

### Statistics in R.

**A note,** If you used the scripts below, we would appreciate that you cite our paper and this publication--Searle, S. R., Speed, F. M., & Milliken, G. A. (1980). Population Marginal Means in the Linear Model: An Alternative to Least Squares Means. The American Statistician, 34(4), 216–221.

Detatils of the version of packages can be found in "Data visualization and statistics", Materials and Methods.

#### Figure 1B and 1C

```
#fig 1B
#Numbers out of the range of mean±2*sd are outliers of the data group.
> c
sample bacterial_growth Days rep
1  Col-0-juv        6.266268 Day2   1
2  Col-0-juv        6.090177 Day2   1
3  Col-0-juv        6.116506 Day2   1
4  Col-0-juv        6.032185 Day2   1
5  Col-0-juv        6.208276 Day2   1
6  Col-0-juv        6.247784 Day2   1
7  Col-0-juv        6.465840 Day2   1
8  Col-0-juv        6.283997 Day2   1
9   fls2-juv        6.477121 Day2   1
10  fls2-juv        5.840299 Day2   1
11  fls2-juv        6.283997 Day2   1
12  fls2-juv        6.164810 Day2   1
```

```
13  fls2-juv         6.454258 Day2    1
14  fls2-juv         6.266268 Day2    1
15  fls2-juv         6.430125 Day2    1
16  fls2-juv         6.090177 Day2    1
17 Col-0-adu         5.477121 Day2    1
18 Col-0-adu         5.442359 Day2    1
19 Col-0-adu         5.317420 Day2    1
20 Col-0-adu         5.718566 Day2    1
21 Col-0-adu         5.712131 Day2    1
22 Col-0-adu         5.301030 Day2    1
23 Col-0-adu         5.585027 Day2    1
24 Col-0-adu         5.649485 Day2    1
25  fls2-adu         6.164810 Day2    1
26  fls2-adu         5.840299 Day2    1
27  fls2-adu         6.090177 Day2    1
28  fls2-adu         6.090177 Day2    1
29  fls2-adu         6.576253 Day2    1
30  fls2-adu         5.789147 Day2    1
31  fls2-adu         6.430125 Day2    1
32 Col-0-juv         2.965238 Day0    1
33 Col-0-juv         3.000000 Day0    1
34 Col-0-juv         3.062148 Day0    1
35  fls2-juv         3.032185 Day0    1
36  fls2-juv         2.927449 Day0    1
37  fls2-juv         2.886057 Day0    1
38 Col-0-adu         3.317420 Day0    1
39 Col-0-adu         3.032185 Day0    1
40 Col-0-adu         3.247784 Day0    1
41  fls2-adu         3.000000 Day0    1
42  fls2-adu         3.164810 Day0    1
43  fls2-adu         3.062148 Day0    1
44  fls2-adu         3.141329 Day0    1
> G=c$bacterial_growth[25:31]
```

```
> G
[1] 6.164810 5.840299 6.090177 6.090177 6.576253 5.789147 6.430125
> df7 <-data.frame(x=4, y0=min(G), y25=mean(G)-2*sd(G), y50=mean(G), y75=mean(G)+2*sd(G), y100=max(G))
> check<-df7$y75
> check
[1] 6.713428
> check<-df7$y25
> check
[1] 5.566854
#fig 1C
> c = read.csv("/Users/lanxi/Desktop/21006_SMii_raw.csv")
> c
samples bacterial_growth Days rep
1    Col_j         3.032185 Day0   1
2    Col_j         2.927449 Day0   1
3    Col_j         2.840299 Day0   1
4    Col_j         2.840299 Day0   1
5   fls2_j         3.333215 Day0   1
6   fls2_j         3.164810 Day0   1
7   fls2_j         3.090177 Day0   1
8   fls2_j         3.090177 Day0   1
9    fec_j         3.062148 Day0   1
10   fec_j         2.927449 Day0   1
11   fec_j         3.062148 Day0   1
12   fec_j         3.032185 Day0   1
13  npr1_j         3.062148 Day0   1
14  npr1_j         3.090177 Day0   1
15  npr1_j         3.247784 Day0   1
16  npr1_j         3.430125 Day0   1
17  npr1_A         3.116506 Day0   1
18  npr1_A         3.090177 Day0   1
19  npr1_A         3.116506 Day0   1
20  npr1_A         3.208276 Day0   1
```

```
21   Col_A          3.090177 Day0   1
22   Col_A          3.164810 Day0   1
23   Col_A          3.208276 Day0   1
24   Col_A          3.090177 Day0   1
25   Col_j          6.404571 Day2   1
26   Col_j          6.567298 Day2   1
27   Col_j          6.283997 Day2   1
28   Col_j          6.641932 Day2   1
29   Col_j          6.090177 Day2   1
30   Col_j          6.032185 Day2   1
31   Col_j          6.477121 Day2   1
32   Col_j          6.247784 Day2   1
33  fls2_j          6.187087 Day2   1
34  fls2_j          6.266268 Day2   1
35  fls2_j          6.465840 Day2   1
36  fls2_j          6.567298 Day2   1
37  fls2_j          6.301030 Day2   1
38  fls2_j          6.228479 Day2   1
39  fls2_j          6.454258 Day2   1
40   fec_j          6.417536 Day2   1
41   fec_j          6.333215 Day2   1
42   fec_j          6.283997 Day2   1
43   fec_j          6.558155 Day2   1
44   fec_j          6.477121 Day2   1
45   fec_j          6.391207 Day2   1
46   fec_j          6.116506 Day2   1
47   fec_j          6.585027 Day2   1
48  npr1_j          7.032185 Day2   1
49  npr1_j          6.927449 Day2   1
50  npr1_j          7.266268 Day2   1
51  npr1_j          6.731155 Day2   1
52  npr1_j          7.348455 Day2   1
53  npr1_j          6.965238 Day2   1
```

```
54  npr1_j          6.927449 Day2   1
55  npr1_j          6.664208 Day2   1
56  npr1_A          6.927449 Day2   1
57  npr1_A          6.927449 Day2   1
58  npr1_A          7.228479 Day2   1
59  npr1_A          7.247784 Day2   1
60  npr1_A          6.840299 Day2   1
61  npr1_A          7.116506 Day2   1
62  npr1_A          6.488117 Day2   1
63  npr1_A          6.363178 Day2   1
64   Col_A          4.731155 Day2   1
65   Col_A          4.927449 Day2   1
66   Col_A          5.141329 Day2   1
67   Col_A          5.116506 Day2   1
68   Col_A          4.927449 Day2   1
69   Col_A          4.927449 Day2   1
70   Col_A          4.488117 Day2   1
> G=c$bacterial_growth[64:70]
> G
[1] 4.731155 4.927449 5.141329 5.116506 4.927449 4.927449 4.488117
> y25=mean(G)-2*sd(G)
> y25
[1] 4.443064
> y75=mean(G)+2*sd(G)
> y75
[1] 5.345352
```

#### Figure 2

```
#fig 2A
> c = read.csv("/Users/lanxi/Desktop/21047_SMii_Juv_D2.csv")
```

```
> c
sample bacterial_growth Days reps
1   Col-0          5.454258 Day2    1
2   Col-0          4.062148 Day2    1
3   Col-0          4.090177 Day2    1
4   Col-0          4.477121 Day2    1
5   Col-0          4.692237 Day2    1
6   Col-0          5.417536 Day2    1
7   Col-0          4.247784 Day2    1
8   Col-0          4.886057 Day2    1
9    fls2          4.626419 Day2    1
10   fls2          4.529509 Day2    1
11   fls2          4.317420 Day2    1
12   fls2          4.266268 Day2    1
13   fls2          4.927449 Day2    1
14   fls2          5.187087 Day2    1
> G=c$bacterial_growth[1:8]
> G
[1] 5.454258 4.062148 4.090177 4.477121 4.692237 5.417536 4.247784 4.886057
> y25=mean(G)-2*sd(G)
> y25
[1] 3.559554
> y75=mean(G)+2*sd(G)
> y75
[1] 5.772275
> G=c$bacterial_growth[9:14]
> G
[1] 4.626419 4.529509 4.317420 4.266268 4.927449 5.187087
> y25=mean(G)-2*sd(G)
> y25
[1] 3.927923
> y75=mean(G)+2*sd(G)
> y75
```

```
[1] 5.356795
#fig 2B
> c = read.csv("/Users/lanxi/Desktop/21054_SMii_Juvd2.csv")
> c
sample bacterial_growth Days reps
1   Col-0         3.164810 Day0    1
2   Col-0         3.164810 Day0    1
3   Col-0         3.404571 Day0    1
4   Col-0         3.404571 Day0    1
5    fls2         2.927449 Day0    1
6    fls2         3.062148 Day0    1
7    fls2         3.247784 Day0    1
8    fls2         3.090177 Day0    1
9   Col-0         7.266268 Day2    1 #out mean±2*sd, an outlier
10  Col-0         6.664208 Day2    1
11  Col-0         6.442359 Day2    1
12  Col-0         6.585027 Day2    1
13  Col-0         6.641932 Day2    1
14  Col-0         6.164810 Day2    1
15  Col-0         6.363178 Day2    1
16  Col-0         6.164810 Day2    1
17   fls2         6.927449 Day2    1
18   fls2         6.363178 Day2    1
19   fls2         6.576253 Day2    1
20   fls2         6.488117 Day2    1
21   fls2         6.333215 Day2    1
22   fls2         7.032185 Day2    1
> G=c$bacterial_growth[9:16]
> G
[1] 7.266268 6.664208 6.442359 6.585027 6.641932 6.164810 6.363178 6.164810
> y25=mean(G)-2*sd(G)
> y25
[1] 5.828437
```

```
> y75=mean(G)+2*sd(G)
> y75
[1] 7.244711
> G=c$bacterial_growth[17:22]
> G
[1] 6.927449 6.363178 6.576253 6.488117 6.333215 7.032185
> y25=mean(G)-2*sd(G)
> y25
[1] 6.032235
> y75=mean(G)+2*sd(G)
> y75
[1] 7.207897
#fig 2C
> c = read.csv("/Users/lanxi/Desktop/22066_smii_hrcCcor_raw.csv")
> c
sample bacterial_growth Days rep
1      Col         2.664208 Day0   1
2      Col         2.363178 Day0   1
3      Col         2.488117 Day0   1
4      Col         2.886057 Day0   1
5     fls2         2.927449 Day0   1
6     fls2         2.488117 Day0   1
7     fls2         2.664208 Day0   1
8     fls2         2.664208 Day0   1
9      Col         5.454258 Day2   1
10     Col         5.247784 Day2   1
11     Col         4.247784 Day2   1
12     Col         4.247784 Day2   1
13     Col         4.417536 Day2   1
14     Col         4.404571 Day2   1
15     Col         5.417536 Day2   1
16     Col         5.141329 Day2   1
17    fls2         5.333215 Day2   1
```

```
18   fls2          4.548814 Day2   1
19   fls2          5.000000 Day2   1
20   fls2          5.247784 Day2   1
21   fls2          4.789147 Day2   1
22   fls2          4.363178 Day2   1
23   fls2          4.488117 Day2   1
24   fls2          4.731155 Day2   1
25    Col          5.794542 Day4   1
26    Col          5.877283 Day4   1
27    Col          5.585027 Day4   1
28    Col          5.558155 Day4   1
29    Col          6.529509 Day4   1
30    Col          7.585027 Day4   1
31   fls2          5.778151 Day4   1
32   fls2          5.576253 Day4   1
33   fls2          5.805135 Day4   1
34   fls2          7.498841 Day4   1
35   fls2          6.825576 Day4   1
36   fls2          5.519525 Day4   1
37   fls2          5.585027 Day4   1
38   fls2          6.164810 Day4   1
> G=c$bacterial_growth[9:16]
> G
[1] 5.454258 5.247784 4.247784 4.247784 4.417536 4.404571 5.417536 5.141329
> y25=mean(G)-2*sd(G)
> y25
[1] 3.743974
> y75=mean(G)+2*sd(G)
> y75
[1] 5.900671
> G=c$bacterial_growth[17:24]
> G
[1] 5.333215 4.548814 5.000000 5.247784 4.789147 4.363178 4.488117 4.731155
```

```
> y25=mean(G)-2*sd(G)
> y25
[1] 4.103721
> y75=mean(G)+2*sd(G)
> y75
[1] 5.521631
> G=c$bacterial_growth[25:30]
> G
[1] 5.794542 5.877283 5.585027 5.558155 6.529509 7.585027
> y25=mean(G)-2*sd(G)
> y25
[1] 4.586897
> y75=mean(G)+2*sd(G)
> y75
[1] 7.72295
> G=c$bacterial_growth[31:38]
> G
[1] 5.778151 5.576253 5.805135 7.498841 6.825576 5.519525 5.585027 6.164810
> y25=mean(G)-2*sd(G)
> y25
[1] 4.669056
> y75=mean(G)+2*sd(G)
> y75
[1] 7.519273
```

#### Figure 4B and 4C

```
#Check outliers for Fig 4B with log2 transformation
#Log2 transformation = Log2(value+1)
> c = read.csv("/Users/lanxi/Desktop/21020_callose_number_raw_log.csv")
> c
```

```
Callose_number rep_names Treatment reps Log2_transformed
1                19   Col-Adu      Mock     1          4.321928
2                 3   Col-Adu      Mock     1          2.000000
3                18   Col-Adu      Mock     1          4.247928
4                22   Col-Adu      Mock     1          4.523562
5                 8   Col-Adu      Mock     1          3.169925
6                31   Col-Adu      Mock     1          5.000000
7                12   Col-Adu      Mock     1          3.700440
8                12   Col-Adu      Mock     1          3.700440
9                57   Col-Adu      Mock     1          5.857981
10              156   Col-Adu     flg22     1          7.294621
11              152   Col-Adu     flg22     1          7.257388
12              155   Col-Adu     flg22     1          7.285402
13               90   Col-Adu     flg22     1          6.507795
14              113   Col-Adu     flg22     1          6.832890
15               95   Col-Adu     flg22     1          6.584963
16               25   Col-Adu     flg22     1          4.700440 #out mean±2*sd, an outlier
17               69   Col-Adu     flg22     1          6.129283
18              118   Col-Adu     flg22     1          6.894818
19               42   Col-Juv     flg22     1          5.426265
20               78   Col-Juv     flg22     1          6.303781
21               11   Col-Juv     flg22     1          3.584963
22               51   Col-Juv     flg22     1          5.700440
23               41   Col-Juv     flg22     1          5.392317
24               43   Col-Juv     flg22     1          5.459432
25               26   Col-Juv     flg22     1          4.754888
26              135   Col-Juv     flg22     1          7.087463
27              160   Col-Juv     flg22     1          7.330917
28                0   Col-Juv      Mock     1          0.000000
29                6   Col-Juv      Mock     1          2.807355
30                5   Col-Juv      Mock     1          2.584963
31                4   Col-Juv      Mock     1          2.321928
32               20   Col-Juv      Mock     1          4.392317
```

```
33     1  Col-Juv   Mock  1   1.000000
34    15  Col-Juv   Mock  1   4.000000
35     5  Col-Juv   Mock  1   2.584963
36    11  Col-Juv   Mock  1   3.584963
37    38 fls2-Adu  flg22  1   5.285402
38    10 fls2-Adu  flg22  1   3.459432
39    37 fls2-Adu  flg22  1   5.247928
40     0 fls2-Adu  flg22  1   0.000000
41     6 fls2-Adu  flg22  1   2.807355
42    10 fls2-Adu  flg22  1   3.459432
43     2 fls2-Adu  flg22  1   1.584963
44     0 fls2-Adu  flg22  1   0.000000
45     0 fls2-Adu  flg22  1   0.000000
46    38 fls2-Adu   Mock  1   5.285402
47    11 fls2-Adu   Mock  1   3.584963
48     0 fls2-Adu   Mock  1   0.000000
49     0 fls2-Adu   Mock  1   0.000000
50     2 fls2-Adu   Mock  1   1.584963
51     0 fls2-Adu   Mock  1   0.000000
52     5 fls2-Adu   Mock  1   2.584963
53    10 fls2-Adu   Mock  1   3.459432
54     3 fls2-Adu   Mock  1   2.000000
55     0 fls2-Juv  flg22  1   0.000000 #out mean±2*sd, an outlier
56    17 fls2-Juv  flg22  1   4.169925
57     5 fls2-Juv  flg22  1   2.584963
58    28 fls2-Juv  flg22  1   4.857981
59    32 fls2-Juv  flg22  1   5.044394
60    27 fls2-Juv  flg22  1   4.807355
61     8 fls2-Juv  flg22  1   3.169925
62    19 fls2-Juv  flg22  1   4.321928
63    13 fls2-Juv  flg22  1   3.807355
64     5 fls2-Juv   Mock  1   2.584963
65     3 fls2-Juv   Mock  1   2.000000
```

```
66              14  fls2-Juv      Mock    1          3.906891
67               2  fls2-Juv      Mock    1          1.584963
68               0  fls2-Juv      Mock    1          0.000000
69              11  fls2-Juv      Mock    1          3.584963
70               0  fls2-Juv      Mock    1          0.000000
71               0  fls2-Juv      Mock    1          0.000000
72               0  fls2-Juv      Mock    1          0.000000
> G=c$Log2_transformed[1:9]
> y25=mean(G)-2*sd(G)
> y25
[1] 1.852403
> y75=mean(G)+2*sd(G)
> y75
[1] 6.263642
> G
[1] 4.321928 2.000000 4.247928 4.523562 3.169925 5.000000 3.700440 3.700440
[9] 5.857981
> G=c$Log2_transformed[10:18]
> y25=mean(G)-2*sd(G)
> y25
[1] 4.972148
> y75=mean(G)+2*sd(G)
> y75
[1] 8.247318
> G
[1] 7.294621 7.257388 7.285402 6.507795 6.832890 6.584963 4.700440 6.129283
[9] 6.894818
> G=c$Log2_transformed[19:27]
> y25=mean(G)-2*sd(G)
> y25
[1] 3.375939
> y75=mean(G)+2*sd(G)
> y75
```

```
[1] 7.966386
> G
[1] 5.426265 6.303781 3.584963 5.700440 5.392317 5.459432 4.754888 7.087463
[9] 7.330917
> G=c$Log2_transformed[37:45]
> y25=mean(G)-2*sd(G)
> y25
[1] -1.861457
> y75=mean(G)+2*sd(G)
> y75
[1] 6.715793
> G
[1] 5.285402 3.459432 5.247928 0.000000 2.807355 3.459432 1.584963 0.000000
[9] 0.000000
> G=c$Log2_transformed[46:54]
> G
[1] 5.285402 3.584963 0.000000 0.000000 1.584963 0.000000 2.584963 3.459432
[9] 2.000000
> y25=mean(G)-2*sd(G)
> y25
[1] -1.680958
> y75=mean(G)+2*sd(G)
> y75
[1] 5.792007
#Emmeans applied for Fig 4C, data of Day2
library(emmeans)
> c = read.csv("/Users/lanxi/Desktop/mean_2sd_20033_SMii_emmeans.csv")
> c
Genotype bacterial_growth Days Treatment rep
1      Col-0          6.116506 Day2      Mock    1
2      Col-0          5.886057 Day2      Mock    1
3      Col-0          6.391207 Day2      Mock    1
4      Col-0          6.488117 Day2      Mock    1
```

```
5      Col-0          6.585027 Day2      Mock   1
6      Col-0          6.465840 Day2      Mock   1
7      Col-0          5.626419 Day2      Mock   1
8      Col-0          5.789147 Day2      Mock   1
9       fls2          6.576253 Day2      Mock   1
10      fls2          6.430125 Day2      Mock   1
11      fls2          6.363178 Day2      Mock   1
12      fls2          6.247784 Day2      Mock   1
13      fls2          5.965238 Day2      Mock   1
14      fls2          6.301030 Day2      Mock   1
15     Col-0          4.965238 Day2     flg22   1
16     Col-0          4.141329 Day2     flg22   1
17     Col-0          4.247784 Day2     flg22   1
18     Col-0          4.965238 Day2     flg22   1
19     Col-0          4.062148 Day2     flg22   1
20     Col-0          4.664208 Day2     flg22   1
21      fls2          5.927449 Day2     flg22   1
22      fls2          5.927449 Day2     flg22   1
23      fls2          5.678448 Day2     flg22   1
24      fls2          6.000000 Day2     flg22   1
25      fls2          5.593627 Day2     flg22   1
> library(emmeans)
> mymodel <- lm(bacterial_growth~Genotype*Treatment, data=c)
> stp1 <- emmeans(mymodel, ~Treatment | Genotype)
> stp2 <- contrast(stp1, "dunnett")
> stp2
Genotype = Col-0:
  contrast     estimate    SE df t.ratio p.value
Mock - flg22     1.661 0.171 21   9.691  <.0001

Genotype = fls2:
  contrast     estimate    SE df t.ratio p.value
Mock - flg22     0.489 0.192 21   2.542  0.0190
```

```
> Genotype_effect <-contrast(stp1, interaction = "pairwise")
> Int.effect <- contrast(Genotype_effect, "dunnett", by=NULL)
> summary(Int.effect)
contrast                                           estimate    SE df t.ratio
(flg22 - Mock fls2) - (flg22 - Mock Col-0)             1.17 0.257 21   4.553
p.value
0.0002
```

#### Figure 6B and 6D

```
#Check outliers for Fig 6B
#Log2 transformation = Log2(value+1)
> c = read.csv("/Users/lanxi/Desktop/21068_SMxix_raw_log2.csv")
> c
    Genotype Number_of_callose Treatment rep Log2_transformation
1        Adu               134     flg22   2            7.076816
2        Adu               119     flg22   2            6.906891
3        Adu               111     flg22   2            6.807355
4        Adu               169     flg22   2            7.409391
5        Adu                78     flg22   2            6.303781
6        Adu               131     flg22   2            7.044394
7        Adu               239     flg22   2            7.906891
8        Adu                98     flg22   2            6.629357
9        Adu               154     flg22   2            7.276124
10       Adu               248     flg22   2            7.960002
11       Adu               202     flg22   2            7.665336
12       Adu               101     flg22   2            6.672425
13       Adu                72     flg22   2            6.189825
14       Adu               104     flg22   2            6.714246
15       Adu                81     flg22   2            6.357552
```

```
16        Adu      152    flg22    2          7.257388
17        Adu      161    flg22    2          7.339850
18        Adu       38    flg22    2          5.285402 #outlier
19        Adu      135    flg22    2          7.087463
20        Adu      160    flg22    2          7.330917
21        Adu      123    flg22    2          6.954196
22        Juv       17    flg22    2          4.169925
23        Juv       30    flg22    2          4.954196
24        Juv       29    flg22    2          4.906891
25        Juv       49    flg22    2          5.643856
26        Juv       39    flg22    2          5.321928
27        Juv       27    flg22    2          4.807355
28        Juv      128    flg22    2          7.011227
29        Juv       35    flg22    2          5.169925
30        Juv      182    flg22    2          7.515700
31        Juv       52    flg22    2          5.727920
32        Juv       28    flg22    2          4.857981
33        Juv       21    flg22    2          4.459432
34        Juv       26    flg22    2          4.754888
35        Juv      135    flg22    2          7.087463
36        Juv       43    flg22    2          5.459432
37        Juv       13    flg22    2          3.807355
38        Juv       49    flg22    2          5.643856
39        Juv        6    flg22    2          2.807355
40        Juv        9    flg22    2          3.321928
41   inMIM156       17    flg22    2          4.169925
42   inMIM156       77    flg22    2          6.285402
43   inMIM156      108    flg22    2          6.768184
44   inMIM156       93    flg22    2          6.554589
45   inMIM156       14    flg22    2          3.906891
46   inMIM156       65    flg22    2          6.044394
47   inMIM156      252    flg22    2          7.982994
48   inMIM156      143    flg22    2          7.169925
```

| | | | | | |
|---|---|---|---|---|---|
| 49 | inMIM156 | 105 | flg22 | 2 | 6.727920 |
| 50 | inMIM156 | 94 | flg22 | 2 | 6.569856 |
| 51 | inMIM156 | 69 | flg22 | 2 | 6.129283 |
| 52 | inMIM156 | 87 | flg22 | 2 | 6.459432 |
| 53 | inMIM156 | 78 | flg22 | 2 | 6.303781 |
| 54 | inMIM156 | 12 | flg22 | 2 | 3.700440 |
| 55 | inMIM156 | 333 | flg22 | 2 | 8.383704 |
| 56 | inMIM156 | 89 | flg22 | 2 | 6.491853 |
| 57 | inMIM156 | 103 | flg22 | 2 | 6.700440 |
| 58 | inMIM156 | 22 | flg22 | 2 | 4.523562 |
| 59 | inMIM156 | 43 | flg22 | 2 | 5.459432 |
| 60 | inMIM156 | 60 | flg22 | 2 | 5.930737 |
| 61 | Adu | 3 | Mock | 2 | 2.000000 |
| 62 | Adu | 2 | Mock | 2 | 1.584963 |
| 63 | Adu | 1 | Mock | 2 | 1.000000 |
| 64 | Adu | 2 | Mock | 2 | 1.584963 |
| 65 | Adu | 39 | Mock | 2 | 5.321928 |
| 66 | Adu | 0 | Mock | 2 | 0.000000 |
| 67 | Adu | 3 | Mock | 2 | 2.000000 |
| 68 | Adu | 0 | Mock | 2 | 0.000000 |
| 69 | Adu | 2 | Mock | 2 | 1.584963 |
| 70 | Adu | 0 | Mock | 2 | 0.000000 |
| 71 | Adu | 3 | Mock | 2 | 2.000000 |
| 72 | Adu | 7 | Mock | 2 | 3.000000 |
| 73 | Adu | 3 | Mock | 2 | 2.000000 |
| 74 | Adu | 3 | Mock | 2 | 2.000000 |
| 75 | Adu | 20 | Mock | 2 | 4.392317 |
| 76 | Adu | 28 | Mock | 2 | 4.857981 |
| 77 | Adu | 9 | Mock | 2 | 3.321928 |
| 78 | Adu | 18 | Mock | 2 | 4.247928 |
| 79 | Adu | 25 | Mock | 2 | 4.700440 |
| 80 | Adu | 35 | Mock | 2 | 5.169925 |
| 81 | inMIM156 | 1 | Mock | 2 | 1.000000 |

```
82  inMIM156      7   Mock  2    3.000000
83  inMIM156     52   Mock  2    5.727920
84  inMIM156     14   Mock  2    3.906891
85  inMIM156     61   Mock  2    5.954196
86  inMIM156      5   Mock  2    2.584963
87  inMIM156     32   Mock  2    5.044394
88  inMIM156     18   Mock  2    4.247928
89  inMIM156     50   Mock  2    5.672425
90  inMIM156     11   Mock  2    3.584963
91  inMIM156      5   Mock  2    2.584963
92  inMIM156      1   Mock  2    1.000000
93  inMIM156     11   Mock  2    3.584963
94  inMIM156      6   Mock  2    2.807355
95  inMIM156     90   Mock  2    6.507795
96  inMIM156     13   Mock  2    3.807355
97  inMIM156     57   Mock  2    5.857981
98  inMIM156     43   Mock  2    5.459432
99       Juv      6   Mock  2    2.807355
100      Juv     15   Mock  2    4.000000
101      Juv     65   Mock  2    6.044394
102      Juv     73   Mock  2    6.209453
103      Juv     10   Mock  2    3.459432
104      Juv     11   Mock  2    3.584963
105      Juv     40   Mock  2    5.357552
106      Juv      9   Mock  2    3.321928
107      Juv     69   Mock  2    6.129283
108      Juv     37   Mock  2    5.247928
109      Juv     30   Mock  2    4.954196
110      Juv     20   Mock  2    4.392317
111      Juv     15   Mock  2    4.000000
112      Juv     14   Mock  2    3.906891
113      Juv     12   Mock  2    3.700440
114      Juv      4   Mock  2    2.321928
```

```
115      Juv                1      Mock    2            1.000000 #outlier
116      Juv               21      Mock    2            4.459432
117      Juv                2      Mock    2            1.584963
118      Juv               22      Mock    2            4.523562
> G=c$Log2_transformation[1:21]
> G
[1] 7.076816 6.906891 6.807355 7.409391 6.303781 7.044394 7.906891
[8] 6.629357 7.276124 7.960002 7.665336 6.672425 6.189825 6.714246
[15] 6.357552 7.257388 7.339850 5.285402 7.087463 7.330917 6.954196
> y25=mean(G)-2*sd(G)
> y25
[1] 5.730061
> y75=mean(G)+2*sd(G)
> y75
[1] 8.191425
> G=c$Log2_transformation[22:40]
> G
[1] 4.169925 4.954196 4.906891 5.643856 5.321928 4.807355 7.011227
[8] 5.169925 7.515700 5.727920 4.857981 4.459432 4.754888 7.087463
[15] 5.459432 3.807355 5.643856 2.807355 3.321928
> y25=mean(G)-2*sd(G)
> y25
[1] 2.711268
> y75=mean(G)+2*sd(G)
> y75
[1] 7.544376
> G=c$Log2_transformation[41:60]
> G
[1] 4.169925 6.285402 6.768184 6.554589 3.906891 6.044394 7.982994
[8] 7.169925 6.727920 6.569856 6.129283 6.459432 6.303781 3.700440
[15] 8.383704 6.491853 6.700440 4.523562 5.459432 5.930737
> y25=mean(G)-2*sd(G)
> y25
```

```
[1] 3.637552
> y75=mean(G)+2*sd(G)
> y75
[1] 8.588723
> G=c$Log2_transformation[61:80]
> y25=mean(G)-2*sd(G)
> y25
[1] -0.9495948
> y75=mean(G)+2*sd(G)
> y75
[1] 6.026328
> G
[1] 2.000000 1.584963 1.000000 1.584963 5.321928 0.000000 2.000000
[8] 0.000000 1.584963 0.000000 2.000000 3.000000 2.000000 2.000000
[15] 4.392317 4.857981 3.321928 4.247928 4.700440 5.169925
> G=c$Log2_transformation[81:98]
> G
[1] 1.000000 3.000000 5.727920 3.906891 5.954196 2.584963 5.044394
[8] 4.247928 5.672425 3.584963 2.584963 1.000000 3.584963 2.807355
[15] 6.507795 3.807355 5.857981 5.459432
> y25=mean(G)-2*sd(G)
> y25
[1] 0.6776565
> y75=mean(G)+2*sd(G)
> y75
[1] 7.359401
> G=c$Log2_transformation[99:118]
> y25=mean(G)-2*sd(G)
> y25
[1] 1.200084
> y75=mean(G)+2*sd(G)
> y75
[1] 6.900517
```

```
> G
[1] 2.807355 4.000000 6.044394 6.209453 3.459432 3.584963 5.357552
[8] 3.321928 6.129283 5.247928 4.954196 4.392317 4.000000 3.906891
[15] 3.700440 2.321928 1.000000 4.459432 1.584963 4.523562
```
